# A multivariate approach to joint testing of main genetic and gene-environment interaction effects

**DOI:** 10.1101/2024.05.06.592645

**Authors:** Saurabh Mishra, Arunabha Majumdar

## Abstract

Gene-environment (GxE) interactions crucially contribute to complex phenotypes. The statistical power of a GxE interaction study is limited mainly due to weak GxE interaction effect sizes. To utilize the individually weak GxE effects to improve the discovery of associated genetic loci, Kraft et al. [1] proposed a joint test of the main genetic and GxE effects for a univariate phenotype. We develop a testing procedure to evaluate combined genetic and GxE effects on a multivariate phenotype to enhance the power by merging pleiotropy in the main genetic and GxE effects. We base the approach on a general linear hypothesis testing framework for a multivariate regression for continuous phenotypes. We implement the generalized estimating equations (GEE) technique under the seemingly unrelated regressions (SUR) setup for binary or mixed phenotypes. We use extensive simulations to show that the test for joint multivariate genetic and GxE effects outperforms the univariate joint test of genetic and GxE effects and the test for multivariate GxE effect concerning power when there is pleiotropy. The test produces a higher power than the test for multivariate main genetic effect for a weak genetic and substantial GxE effect. For more prominent genetic effects, the latter performs better with a limited increase in power. Overall, the multivariate joint approach offers high power across diverse simulation scenarios. We apply the methods to lipid phenotypes with sleep duration as an environmental factor in the UK Biobank. The proposed approach identified six independent associated genetic loci missed by other competing methods.

## 1 Introduction

Epidemiological studies have recognized relationships between environmental factors and disease risk or phenotype variation. However, all individuals exposed to a specific environmental factor, e.g., smoking, do not develop a disease, e.g., lung cancer. Similarly, not all individuals who inherit particular genetic variants develop a disease. Thus, combinations of genetic and environmental factors, hence gene by environment (GxE) interactions, play a paramount role in many disease phenotypes [2]. Characterizing GxE interactions helps to understand how environmental factors can modulate genetic predisposition to a complex phenotype [3, 4]. Genome-wide association study (GWAS) is the standard tool for discovering phenotype-associated genetic loci. Similarly, genome-wide genetic variants, such as SNPs, are tested for the GxE interaction effect between a phenotype and environmental factor [5].

For example, Maazi et al. [6] identified GxE interaction between diesel exhaust-elicited airway hyper-reactivity and a genetic locus on chromosome 3, which encodes the gene DAPP1. Pierce et al. [7] found that rs9527 on chromosome 10 interacts with arsenic exposure to influence the risk of skin lesions in the Bangladeshi population. Despite some exciting discoveries, GxE interaction studies are yet to see abundant success due to inadequate statistical power. Instead of a single variant level test for GxE interaction, some recent studies focused on characterizing aggregate genome-wide GxE polygenic contribution to a complex phenotype [8]. While such studies provide important insights into the puzzle of missing heritability and phenotype biology, they generally lack the resolution of variant-level investigations of GxE interactions. To improve the power of variant-level GxE interaction tests, various alternative approaches, such as case-only studies under the gene-environment independence assumption [9, 10, 11], two-step approaches that filter out less critical variants in an initial step to reduce multiple-testing burden [12, 13, 4, 14, 15], etc., have been proposed. However, such methods are yet to successfully detect numerous reproducible GxE interactions as hoped [16, 17]. These findings indicate that GxE interaction effects are minor, and we require substantial sample sizes to detect them reliably.

Instead of discovering the GxE interaction effect alone, Kraft et al. [1] proposed that searching for a joint effect of marginal genetic or GxE interaction factors can boost the power to find the genetic loci associated with the phenotype. For a disease phenotype, a locus can modify the risk only for those individuals exposed to an environmental factor. Here, the locus has a GxE interaction but no marginal genetic effect. When the locus has a marginal effect but no interaction effect, it modulates the risk irrespective of the exposure status. Since the actual mechanism is unknown, a joint test for GxE interaction and marginal genetic effects is more robust and consistently powerful [1, 13]. In this manner, the individually weak GxE effects can complement the main genetic effects to locate the susceptibility loci for the phenotype better. The joint test is powerful, especially for variants with minor to moderate main and interaction effects [18].

Multi-phenotype approaches analyze the genetic architecture of multiple related phenotypes simultaneously instead of a single phenotype because genetic effects are often shared across phenotypes (pleiotropy), e.g., lipid phenotypes, LDL, HDL, and triglycerides. Jointly modeling multiple phenotypes increases statistical power, uncovers pleiotropic effects, and provides a more comprehensive understanding of their genetic architecture [19, 20, 21, 22, 23]. Not only for the main genetic effect but Majumdar et al. [15] also demonstrated that modeling pleiotropy in GxE effects produces better power than testing for a univariate GxE effect. We hypothesize that considering a multivariate phenotype while jointly testing for the main genetic or GxE interaction effect can combine the advantage of the two domains, integrated modeling of the main genetic and GxE effect, and pleiotropy.

In this article, we develop novel approaches to joint testing of the main genetic and GxE interaction effects for a multivariate phenotype, which can be continuous, binary, or a mixture of both. We employ the procedure for testing a general linear hypothesis in a multivariate regression for continuous phenotypes. We implement the generalized estimating equations (GEE) technique in a seemingly unrelated regressions (SUR) framework when the phenotypes are binary or have mixed types. Using extensive simulations, we compare the power of competing alternatives – tests for joint multivariate genetic and GxE effects, joint univariate genetic and GxE effects, multivariate GxE effects, multivariate genetic effects, etc. The combined test for multivariate genetic and GxE effects performs better in most simulation settings and produces robustly high power across diverse scenarios. We apply the methods to UK Biobank data [24] for genome-wide screening of GxE effects for lipid phenotypes, LDL, HDL, and triglycerides while considering sleep duration as the environmental factor. Our multivariate joint approach identified six independent associated loci that other methods overlooked.

## 2 Methods

Consider phenotype and genotype data for *n* unrelated individuals. Suppose the multivariate phenotype comprises *m* phenotypes, *Y* = (*Y*_1_, *…, Y*_*m*_), either continuous or binary or a mixture of both. Consider an SNP for which the genotype data is available for the individuals. Genotype is coded as the number of minor alleles in the genotype, and *G* denotes the coded value. Let *E* denote an environmental factor. We first consider the case when all the phenotypes are continuous. We use multivariate multiple linear regression (MMLR) [25] to model the relationship between the phenotype vector and the genotype, environmental factor, and their interaction. MMLR can efficiently model a covariance structure of the phenotypes. Next, we outline the regression frameworks and hypotheses to test for various combinations of the main genetic and GxE interaction effects in the multivariate setup. We begin with the existing approaches to individually testing a multivariate genetic effect and GxE effect. Then, we describe the layout for our joint multivariate approach. Finally, we also provide the univariate set-up used for comparative evaluation in the paper.

### 2.1 Multivariate main genetic effect

We consider the following multivariate linear regression to test for an overall main genetic effect of the SNP on the multivariate phenotype:

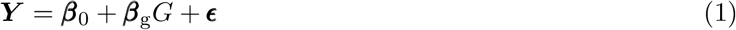

Here, ***β***_0_ = (*β*_01_, *…, β*_0*m*_)^′^ is the intercept vector, and ***β***_g_ = (*β*_g1_, *…, β*_g*m*_)^′^ denotes the vector of main genetic effects of the SNP on *Y*. We test for 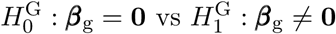 to test for an overall pleiotropy in main genetic effects. ***β***_g_≠ 0 implies that *β*_g*j*_≠ 0 for at least one *j* = 1, *…, m*. The vector of residual components, **ϵ** = (ϵ_1_, · · ·, ϵ_*m*_), is assumed to follow multivariate normal: **ϵ** ∼ *N*_*m*_(0, **Σ**).

### 2.2 Multivariate GxE interaction effect

We regard the following MMLR to test for an overall GxE interaction effect. The MMLR framework offers great flexibility in testing various combinations of hypotheses on the regression coefficients across phenotypes.

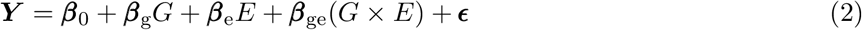

*G*×*E* denotes the interaction term. ***β***_g_ = (*β*_g1_, *…, β*_g*m*_)^′^, ***β***_e_ = (*β*_e1_, *…, β*_e*m*_)^′^, and ***β***_ge_ = (*β*_ge1_, *…, β*_ge*m*_)^′^ denote the vectors of main genetic, environmental, and GxE interaction effects on *Y*, respectively. To test for an overall pleiotropy in the GxE effect, we test for 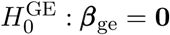 versus 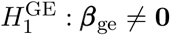 While testing 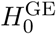, we compare the unrestricted model (equation 2) with the reduced model Y = ***β***_0_ + ***β***_g_*G* + ***β***_e_*E +* ***ϵ*** under 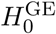 using the type II MANOVA.

### 2.3 Joint multivariate main genetic and GxE interaction effects

We consider the following hypothesis under the MMLR model (equation 2) to test for a joint multivariate main genetic and GxE interaction effects, 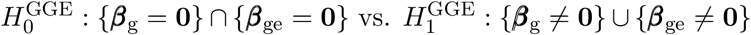 Under 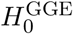, there are no multivariate effects due to the genetic and GxE factors. There are three possibilities under the alternate hypothesis: non-zero main genetic but null GxE interaction effect, null main genetic but non-zero GxE interaction effect, and non-zero main genetic and GxE interaction effects. As mentioned above, a non-zero multivariate effect implies that at least one element of the effect size vector is non-zero. To test for 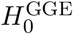, we again compare the unrestricted model (equation 2) with the reduced model Y = ***β***_0_ + ***β***_e_*E* + ***ϵ*** under 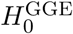using the type II MANOVA.

### 2.4 Univariate set-up

We employ the multiple linear regression model for univariate analysis of *Y*_*j*_, *j* = 1, *…, m*. To test for genetic effect, consider *Y*_*j*_ = *β*_0*j*_ + *β*_g*j*_ *G* + ϵ_*j*_, and for GxE effect, consider *Y*_*j*_ = *β*_0*j*_ + *β*_g*j*_ *G* + *β*_e*j*_ *E* + *β*_ge*j*_ (*G* × *E*) + ϵ_*j*_. Here, *β*_g*j*_, *β*_e*j*_ and *β*_ge*j*_ denote univariate genetic, environmental, and GxE effects for *j*^*th*^ phenotype, respectively; and the residual term 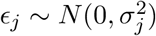, for *j* = 1, *…, m*. To evaluate the univariate genetic effect for *Y*_*j*_, test for 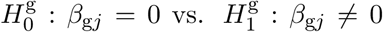 To assess the univariate GxE effect, test for 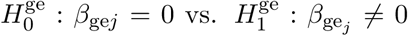. To evaluate the univariate joint genetic and GxE effects, test for 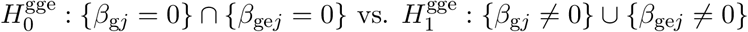

To summarize, we consider the following three pairs of null and alternative hypotheses in the multi-variate scenarios:

1. To test for multivariate main genetic effect – 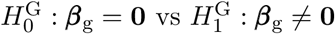
2. To test for multivariate GxE interaction effect – 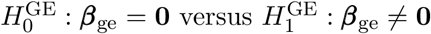
3. To test for joint multivariate main genetic and GxE interaction effects – 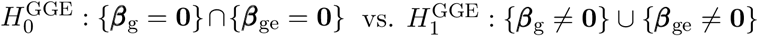

### 2.5 Hypothesis testing

Consider the following MMLR in matrix notation for *n* unrelated individuals’ data:

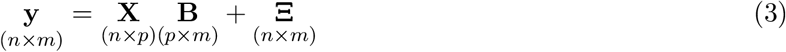

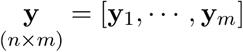 denotes the matrix of phenotype values for *n* individuals and *m* phenotypes such that y_*j*_ = (*y*_1*j*_, · · ·, *y*_*nj*_)^′^ is the vector of *j*^*th*^ phenotype values, 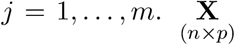 is the design matrix with columns for *p* regressors, including the first column of 1s corresponding to the intercept coefficient. 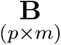 is the matrix of regression coefficients, with one column for each phenotype. 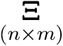 is the matrix of error terms with columns corresponding to the phenotypes. Our goal is to test a general linear hypothesis for the regression coefficients in the MMLR model [25, 26]:

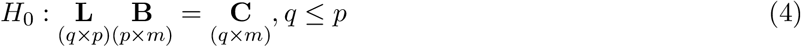

Here, 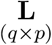 is the hypothesis matrix with full row-rank, which imposes *q* linearly independent constraints on 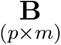, the matrix of regression coefficients. 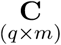 matrix consists of relevant constants, usually zero column vectors for the target null hypothesis. To test for a specific combination of multivariate genetic and GxE effects, we can choose the L and C matrices accordingly in equation 4 to obtain the target *H*_0_.

#### 2.5.1 Choices of L and C matrices

We regard 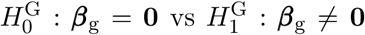 to test the multivariate genetic effect. Here, the B matrix of regression coefficients takes the following form for *m* phenotypes:

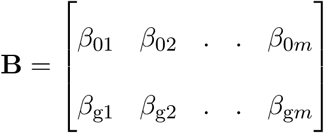

The first entry of each column representing a phenotype denotes the intercept coefficient. Choosing 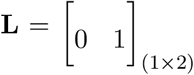 and 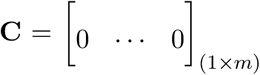 in equation 4 induces *H*_0_ : (*β*_g1_, …, *β*_g*m*_) = (0, …, 0); applying a transpose to both sides of the equation for *H*_0_ provides the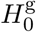.

To test for multivariate GxE effect, we consider 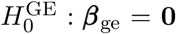 versus 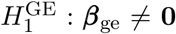 Regard the B matrix as:

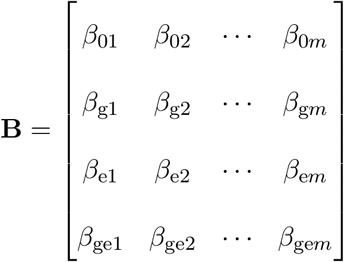

Choosing 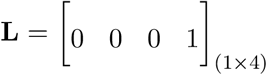 and 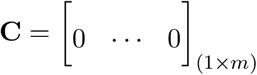 results in the target null hypothesis 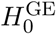

Next, to test for a joint multivariate genetic and GxE effects, we evaluate 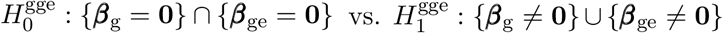 Consider the above B matrix that includes all the coefficients for genetic, environmental, and GxE effects. Choosing 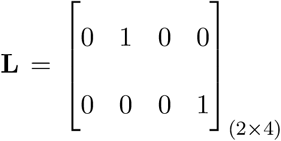 and 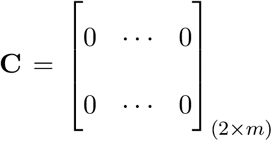 leads to 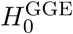

We use Wilk’s Lambda test statistic under the type II MANOVA framework to test these three hypotheses. We implement the R package ‘*car*’ [27] to perform the type II MANOVA. More details about Wilk’s Lambda test statistic are provided in the supplementary materials.

### 2.6 Binary or mixed phenotypes

A multivariate phenotype can constitute all binary or mixed phenotypes, such as one binary and one continuous, e.g., type 1 diabetes and LDL cholesterol. Modeling a multivariate binary or mixed phenotype is more challenging than the continuous case. For simplicity, we describe the methods for a bivariate mixed or binary phenotype (*m* = 2), which can be extended for more phenotypes. In these scenarios, we adapt the generalized estimating equations (GEE) technique [28, 29] in seemingly unrelated regressions (SUR) framework [30].

#### 2.6.1 Bivariate mixed phenotype

We first consider a bivariate mixed phenotype comprising one continuous and one binary phenotype. Suppose we have data on two correlated phenotypes *Y*_1_ and *Y*_2_ such that *Y*_1_ is normally distributed, and *Y*_2_ is binary. Let *y*_*i*_ = (*y*_1*i*_, *y*_2*i*_)^′^ denote the phenotype vector for *i*^*th*^ individual with the corresponding mean vector, *µ*_*i*_ = (*µ*_1*i*_, *µ*_2*i*_)^′^ and the covariance matrix, *V* _*i*_ = cov(*y*_*i*_), *i* = 1, · · ·, *n*. Let *x*_*i*_ denote *p*-length vector of explanatory variables for *i*^*th*^ individual, e.g., the genotype, environmental exposure, and GxE term. Let ***β***_1_ and ***β***_2_ denote *p*-length vectors of regression coefficients corresponding to *Y*_1_ and *Y*_2_, respectively. We employ the generalized linear model (GLM) to express the relationship between the explanatory variables and the marginal means of *Y*_1_ and *Y*_2_ as:

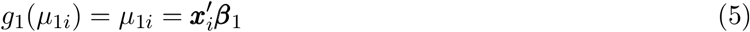

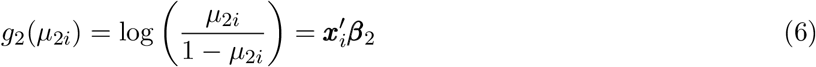

Here, *g*_1_(.) and *g*_2_(.) are the identity and logit link functions for *Y*_1_ and *Y*_2_, respectively. We need to integrate the above two phenotype-specific models into a unified structure to account for the covariance between the phenotypes. We implement the seemingly unrelated regression (SUR) framework [30] to combine the individual relationships for the two phenotypes and estimate the parameters of the regression models. Based on the SUR setup,

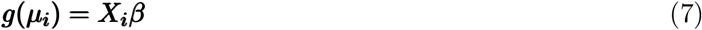

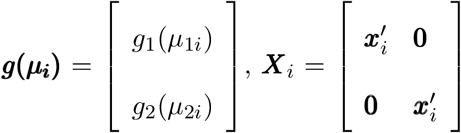 and 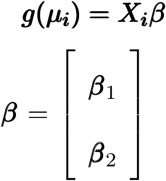 The dimesions of *g*(*µ*_*i*_)*X*_*i*_, and *β* are 2×1,2 × 2*p*, and 2*p* × 1, respectively.

Using GLM theory, let *v*_1*i*_ = var (*y*_1*i*_) = *ψ*_1_ ν_1_ (*µ*_1*i*_) and *v*_2*i*_ = var (*y*_2*i*_) = *ψ*_2_ ν_2_ (*µ*_2*i*_), where ν_1_(·) and ν_2_(·) are known variance functions. *ψ*_1_ and *ψ*_2_ are dispersion parameters for normal and binomial distributions, respectively. For binary *Y*_2_, *ψ*_2_ = 1 as there is no overdispersion. For *Y*_1_, we consider a squared transformation *ψ*_1_ = *φ*^2^ for convenience in derivations, where *ψ*_1_ *>* 0. The variance functions are specified as:

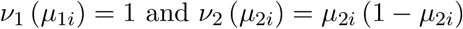

To model the correlation structure of the mixed phenotypes, we consider the generalized estimating equations (GEE) approach [28, 29], which specifies a working correlation structure of the phenotypes. The GEE technique performs reliably even if the correlation structure is not completely accurate. The method treats the correlation parameters as nuisance parameters in the estimation process [28, 31]. The working correlation matrix can have different structures depending on the assumptions of the correlation pattern, such as independent, exchangeable, autoregressive, etc. We consider an exchangeable correlation pattern for *Y*_*1*_, *Y*_*2*_. Let 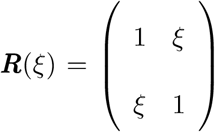 be the working correlation matrix, where ξ is an unknown association parameter taking values in (−1,1). *R*_*i*_(ξ), the working correlation structure for *i*^*th*^ individual is chosen to be the same as *R*(ξ), *i* = 1, · · ·, *n*. Based on the choices of the working correlation matrix, variance functions, and dispersion parameters, we obtain the covariance matrix of (*Y*_1_, *Y*_2_) for *i*^*th*^ individual as:

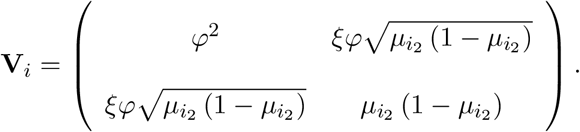

Next, we outline the GEE estimation procedure. Let **α** = (ξ, *φ*)^′^ denote the vector of second-order moment parameters. A solution to the following generalized estimating equations provides the GEE estimator of ***β***.

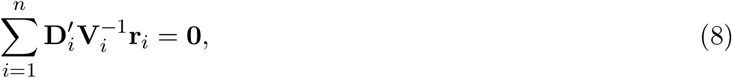

where ***r*_*i*_ = *y*_*i*_ − *µ*_*i*_**, and

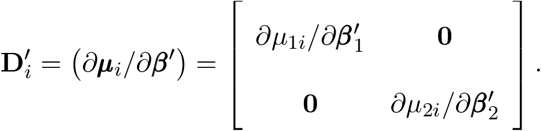

Given that **D**_*i*_, **V**_*i*_, and **r**_*i*_, *i* = 1, …, *n*, are evaluated at a consistent estimator of **α**, Liang and Zeger [28, 29] demonstrated that the estimated vector of regression coefficients 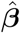 asymptotically follows a normal distribution. Furthermore, 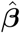 is asymptotically unbiased with the following asymptotic covariance matrix:

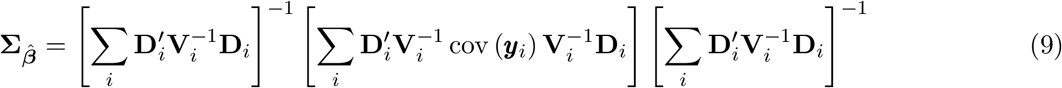

This quantity can be estimated by evaluating the matrices in the expression at the GEE estimates of the unknown parameters and replacing cov (*y*_*i*_) by 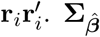 is a consistent estimator even if V_*i*_ is not specified correctly. A robust estimator 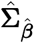 for the covariance matrix can be obtained by replacing *cov*(***y*_*i*_**) with 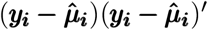, where 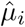 is evaluated at 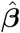.

An overall advantage of the GEE method is that the large sample properties of 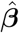and 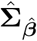 hold even if an utterly correct correlation matrix among the observations can not be specified. However, those correlation matrices closer to the truth will offer a comparatively better efficiency. Nevertheless, Zeger [32] concluded that the gain in efficiency becomes negligible for a large sample size, which is expected for a contemporary large-scale genetic study. Since there is no closed-form solution to the set of equations (equation 8), Liang and Zeger [28] described an iterative scoring algorithm to obtain 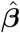. Given the parameters*ψ* estimates at *l*^*th*^ iteration, **α**^(*l*+1)^ is obtained by moment estimators. Given **α**^(*l*+1)^, *β*^(*l*+1)^ is obtained by updating *β*^(*l*)^ according to the following scoring step:

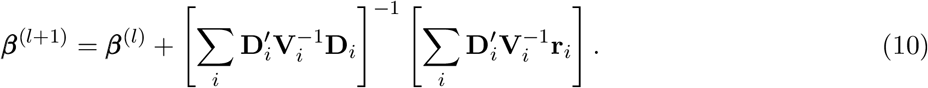

More details of the estimation algorithm are provided in the supplementary materials.

Estimated 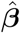provides an estimate of the B matrix of regression coefficients for general linear hypothesis testing (equation 4). The asymptotic covariance matrix for 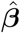(equation 9) provides an estimate of the covariance structure of B. For a given choice of *H*_0_ and the corresponding L and C matrices in equation 4, we use the Wald’s test statistic to test *H*_0_:

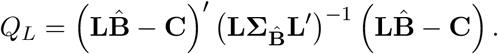

*Q*_*L*_ asymptotically follows *χ*^2^ with degree of freedom *q*, where *q* is the number of rows in the L matrix. The choice of *H*_0_ depends on the combination of the multivariate genetic and GxE effects to be tested for.

#### 2.6.2 Bivariate binary phenotype

When the bivariate phenotype comprises binary phenotypes, the models, estimation, and hypothesis testing procedure remain similar to the mixed phenotype case. The main difference is in the model specification under the GLM framework, where we replace the continuous with a binary phenotype and the identity link function with a logit link function (equation 5).

## 3 Simulation Study

We perform extensive simulations to compare our approach concerning the type I error rate and power with the competing approaches. For a multivariate phenotype, we compare the test for a combined multivariate genetic and GxE effects with the test for a joint univariate genetic and GxE effect on each phenotype. We also explore the performance of the competing approaches that individually test for a multivariate genetic effect and GxE effect.

### 3.1 Setup

We simulate the unrelated individuals*ψ* genotypes at an SNP and environmental factor data. We use simulation models to generate the phenotype data based on the genotype and environmental factor measurements. We assume that the SNP follows the Hardy-Weinberg equilibrium (HWE). Let *A, a* denote the alleles for the SNP. Under HWE, the genotype probabilities are *P* (*AA*) = *p*^2^, *P* (*Aa*) = 2*p*(1 − *p*), and *P* (*aa*) = (1 − *p*)^2^, where *P* (*A*) = *p*. We simulate a binary environmental exposure *E* with *P* (*E* = 1) = *f* = 1 − *P* (*E* = 0). For simplicity, we consider a bivariate phenotype in the simulations. Let (*Y*_1_, *Y*_2_)′ denote the bivariate phenotype, *G* denote the genotype at the SNP. We consider the following bivariate linear regression model to simulate the bivariate continuous phenotype.

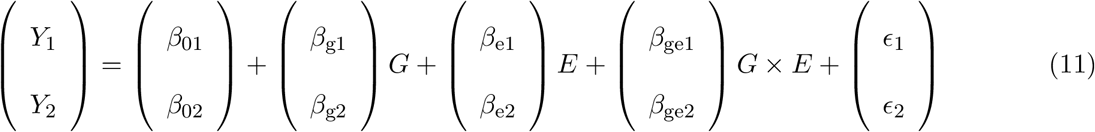

We assume that (ϵ_1_, ϵ_2_) follows bivariate normal with the means as zero and the variances as 0.5 and the correlation as 0.3. We choose a sample size of 1000. We select the minor allele frequency (MAF) for the SNP as 0.1 and the proportion of the reference category for the environmental exposure as *f* = 0.1. We select the effect sizes of the environmental factor as *β*_e1_ = *β*_e2_ = *β*_e_ = 0.2. We only consider a non-zero value of *β*_e_ because we assume that environmental exposure always affects the phenotypes regardless of the genetic and GxE effects. We assume the same genetic effects for the phenotypes, *β*_g1_ = *β*_g2_ = *β*_g_, and let *β*_g_ take a value in {0, 0.05, 0.1, 0.15, 0.2}. Similarly, the GxE effects are the same, *β*_ge1_ = *β*_ge2_ = *β*_ge_, and *β*_ge_ ∈ {0, 0.1, 0.2, 0.3, 0.4, 0.5}. We assume equal effect sizes for both phenotypes to ease the description of results. Thus, when *β*_g_ or *β*_ge_ is non-zero, there is pleiotropy in the genetic or GxE effects. To evaluate the powers when there is no pleiotropy in the genetic and GxE effects, we assume the second phenotype-specific effects to be zero, *β*_g2_ = *β*_ge2_ = 0. We regard *β*_g_ = *β*_ge_ = 0 while simulating phenotype data to evaluate the false positive rates of the methods under various choices of the null hypothesis. We conduct 1000 iterations to estimate various tests’ type I error rate and power for a given simulation scenario.

We apply thresholding on *Y*_1_ after generating (*Y*_1_, *Y*_2_) as described above to generate a bivariate mixed phenotype. We choose the threshold *t* as the 70^*th*^ sample percentile of *Y*_1_. Next, define the binary phenotype *Z*_1_ as: *Z*_1_ = 1 if *Y*_1_ *> t* and *Z*_1_ = 0 otherwise. We regard (*Z*_1_, *Y*_2_) as the bivariate mixed phenotype. Thus, for a constituent binary phenotype in the bivariate phenotype, 30% of the individuals are cases, and 70% are controls. Similarly, we generate the bivariate binary phenotype by thresholding *Y*_1_ and *Y*_2_.

We perform univariate and multivariate tests for the genetic, GxE, and combined genetic and GxE effects. Previous studies compared the power between univariate and multivariate approaches to individually testing genetic [33, 20] and GxE effects [15]. These studies demonstrated higher power for the multivariate approaches while individually testing for the genetic and GxE effects. For a concise presentation, we compare our approach to testing combined multivariate genetic and GxE effects with the two degrees of freedom univariate joint test of genetic and GxE effects proposed by Kraft et al. [1]. We include separate tests for multivariate genetic and GxE effects as competing approaches. We denote the multivariate joint test as the *GGE* test and the univariate joint test as the *gge* test. Individual tests for multivariate genetic and GxE effects are termed *G* and *GE* tests, respectively.

### 3.2 Simulation results

All methods control the type I error rates while testing various null hypotheses of genetic, GxE, and joint genetic and GxE effects for univariate and multivariate phenotypes (Table 1). The results are consistent across continuous, mixed, and binary phenotypes. We sometimes observe marginal inflations and deflations in the estimated rates (Table 1), which can be attributed to sampling fluctuations.

**Table 1:** Estimated type I error rates for different multivariate tests for various choices of the null hypotheses. The *G* test evaluates the multivariate genetic effect. The *GE* test evaluates the multivariate GxE effect. The *GGE* test assesses the combined multivariate genetic and GxE effects.

|  |  | Tests for bivariate |  |  | Tests for bivariate |  |  | Tests for bivariate |  |  |
| --- | --- | --- | --- | --- | --- | --- | --- | --- | --- | --- |
|  |  | continuous phenotypes |  |  | mixed phenotypes |  |  | binary phenotypes |  |  |
| MAF | $P(E = 1)$ | $G$ | $GE$ | $GGE$ | $G$ | $GE$ | $GGE$ | $G$ | $GE$ | $GGE$ |
| 0.1 | 0.1 | 0.05 | 0.05 | 0.05 | 0.04 | 0.07 | 0.06 | 0.04 | 0.05 | 0.05 |
| 0.1 | 0.25 | 0.05 | 0.05 | 0.05 | 0.05 | 0.05 | 0.06 | 0.06 | 0.04 | 0.04 |
| 0.25 | 0.1 | 0.04 | 0.06 | 0.05 | 0.04 | 0.06 | 0.06 | 0.04 | 0.06 | 0.06 |
| 0.25 | 0.25 | 0.04 | 0.04 | 0.04 | 0.06 | 0.05 | 0.05 | 0.05 | 0.05 | 0.05 |

Concerning power, the *GGE* test performs consistently better than the two degrees of freedom *gge* test (Figures 1, 2, 3) when there is pleiotropy in the genetic and GxE effects. In these simulation scenarios, the power increase of the *GGE* test than the *gge* test varies in the range of 1 − 20% for a bivariate continuous phenotype (Figure 1), 3 − 14% for a bivariate binary phenotype (Figure 2), and 2 − 18% for a mixed phenotype (Figure 3). Here, we regard the maximum power of the *gge* test across the phenotypes. When there is no pleiotropy in the genetic and GxE effects, the *gge* test for the first phenotype performs better than the *GGE* test (Figure 4). The *gge* test has almost no power for the second phenotype as there is no genetic and GxE effect on the phenotype (Figure 4). Since there is no pleiotropy, the *GGE* test does not gain any benefit but uses two extra degrees of freedom. However, the power loss of the *GGE* test compared to *gge* test for the first phenotype is limited, varying in 1 − 8% when the genetic effect for the first phenotype is small to moderate (Figure 4). For a more potent genetic effect (*β*_g1_ = 0.2), the powers between the two approaches become comparable (Figure 4).

**Figure 1.**
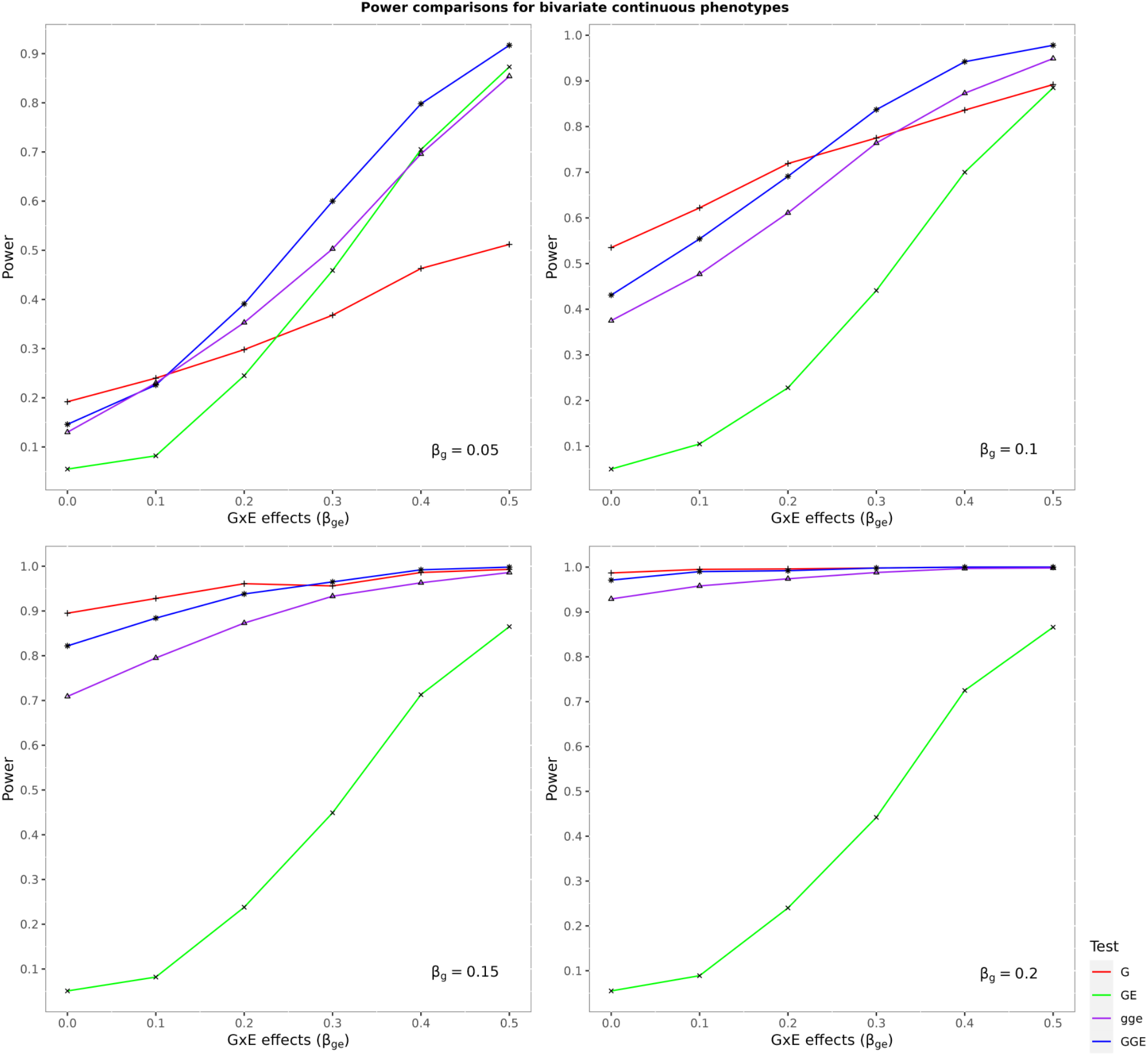
Comparison of powers obtained by different tests for bivariate continuous phenotypes in the presence of pleiotropy. The *G* test evaluates the multivariate genetic effect. The *GE* test evaluates the multivariate GxE effect. The *GGE* test assesses the combined multivariate genetic and GxE effects. The *gge* test gauges the joint univariate genetic and GxE effects. Since the genetic and GxE effect sizes are the same for both simulated phenotypes, the power curve for the *gge* test is based on the maximum power obtained across the univariate phenotypes.

**Figure 2.**
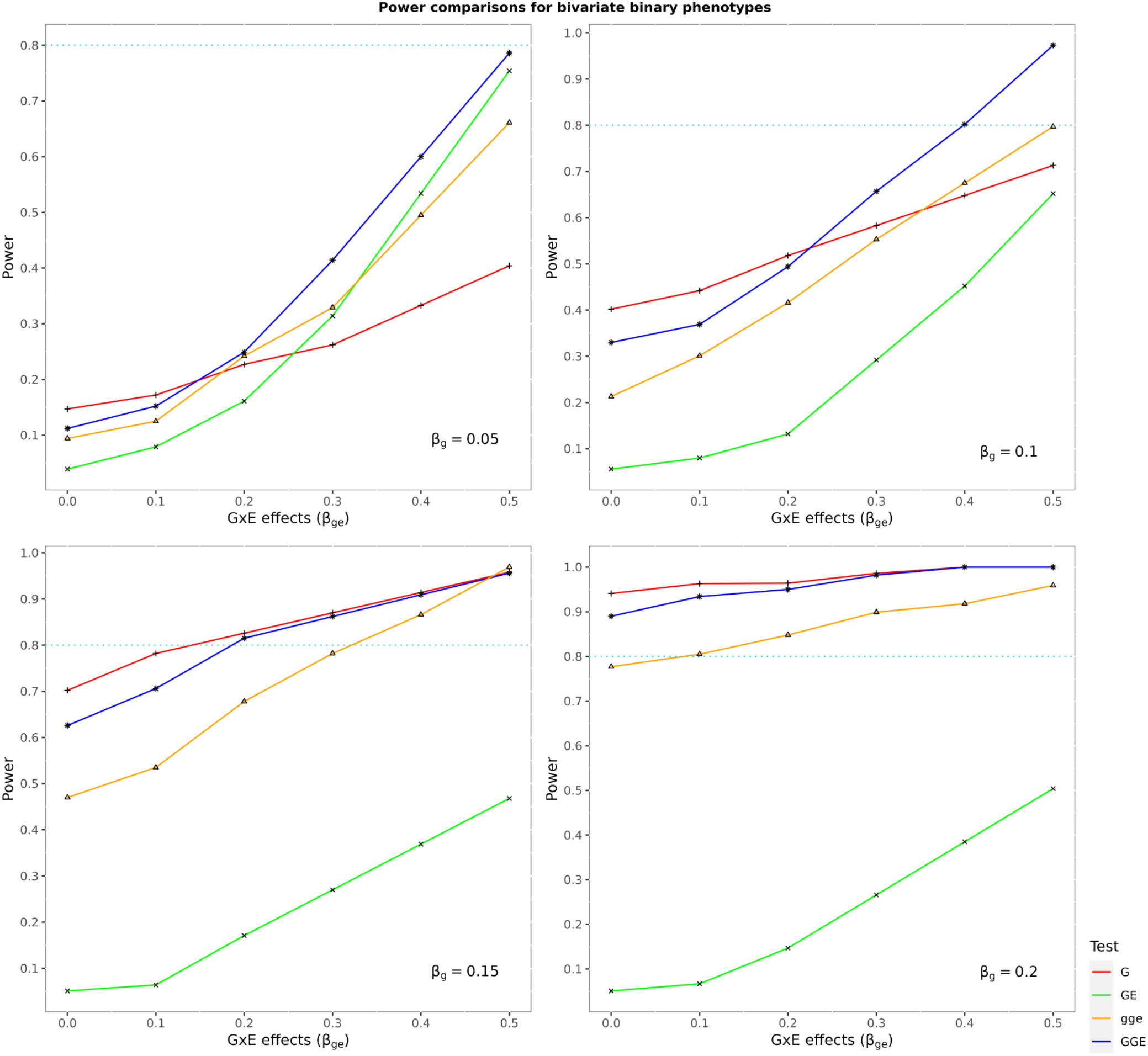
Comparison of powers obtained by different tests for bivariate binary phenotypes in the presence of pleiotropy. The *G* test evaluates the multivariate genetic effect. The *GE* test evaluates the multivariate GxE effect. The *GGE* test assesses the combined multivariate genetic and GxE effects. The *gge* test gauges the joint univariate genetic and GxE effects. Since the genetic and GxE effect sizes are the same for both simulated phenotypes, the power curve for the *gge* test is based on the maximum power obtained across the univariate phenotypes.

**Figure 3.**
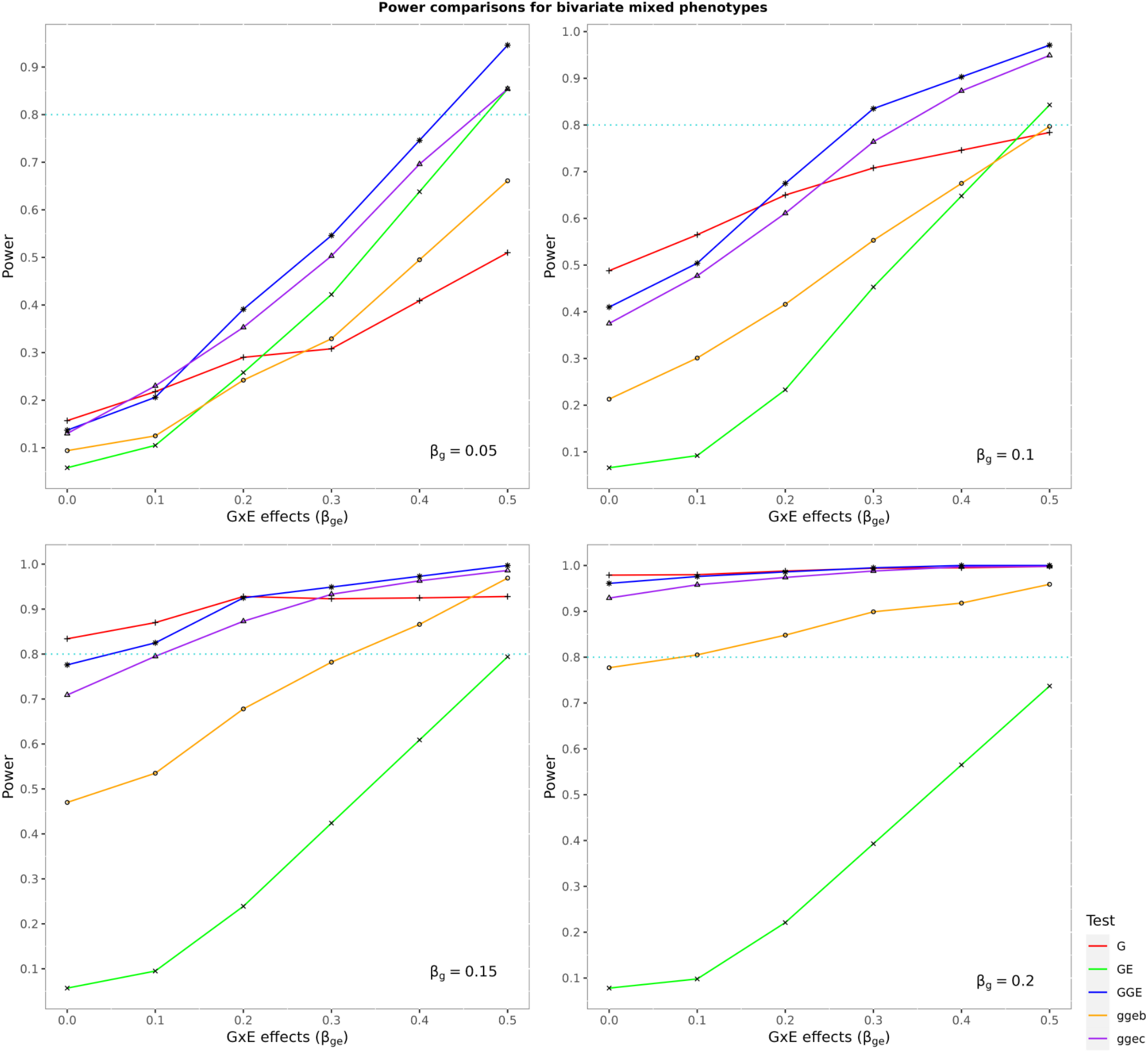
Comparison of powers obtained by different tests for bivariate mixed phenotypes in the presence of pleiotropy. The *G* test evaluates the multivariate genetic effect. The *GE* test evaluates the multivariate GxE effect. The *GGE* test assesses the combined multivariate genetic and GxE effects. The *gge*c and *gge*b tests gauge the joint univariate genetic and GxE effects for the continuous and binary phenotypes, respectively.

**Figure 4.**
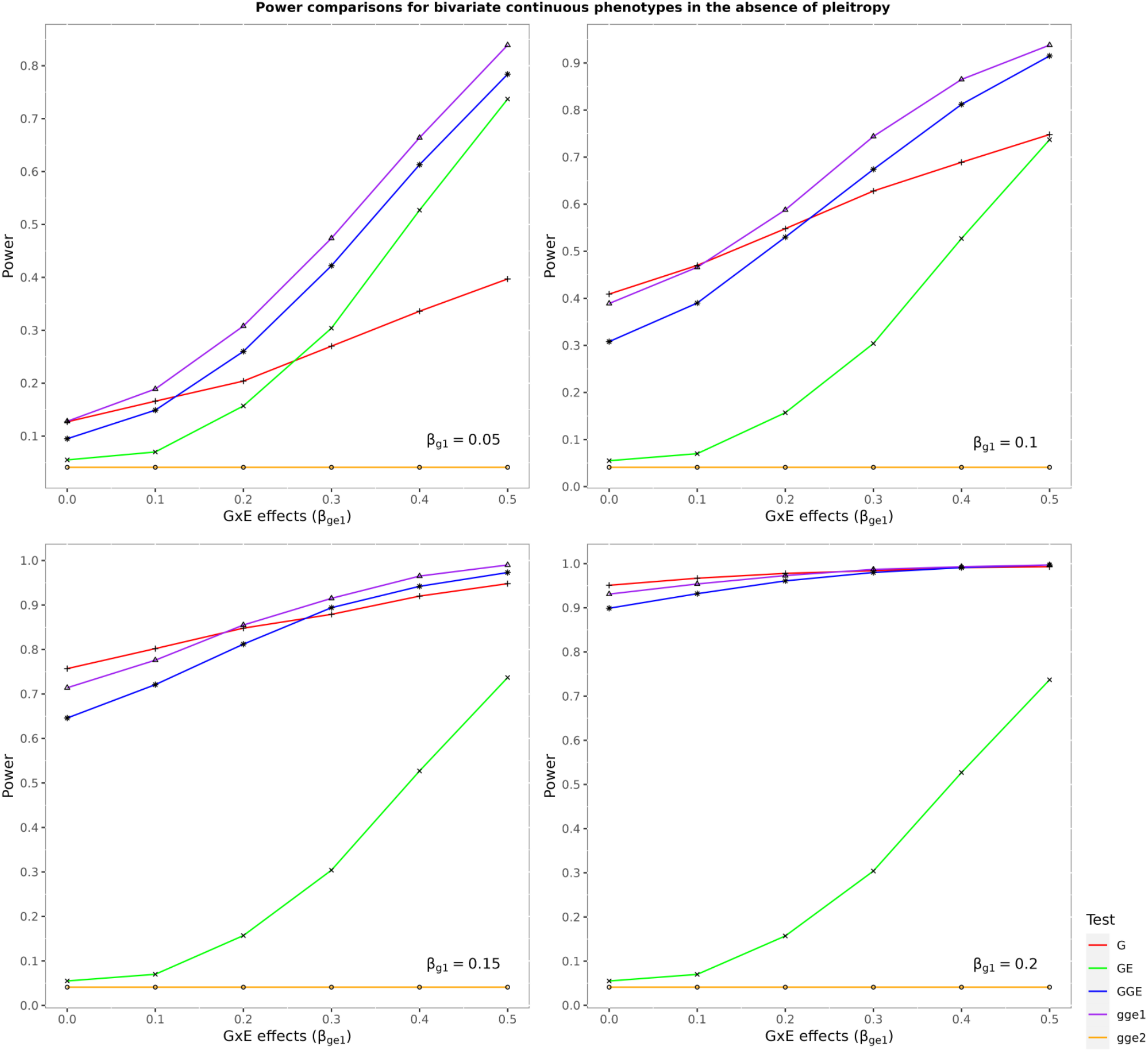
Comparison of powers obtained by different tests for bivariate continuous phenotypes without pleiotropy. The *G* test evaluates the multivariate genetic effect. The *GE* test evaluates the multivariate GxE effect. The *GGE* test assesses the combined multivariate genetic and GxE effects. The *gge*1 and *gge*2 tests gauge the joint univariate genetic and GxE effects for the first and second phenotypes, respectively.

Next, we compare the performance of the *GGE* test and the *GE* test for multivariate GxE effect. The *GGE* test outperforms the *GE* test with a power increase of 3 − 90% for continuous phenotypes (Figure 1), 3 − 86% for binary phenotypes (Figure 2), and 7 − 87% for mixed phenotypes (Figure 3). The former results in a power increase of 4 − 86% than the latter when there is no pleiotropy (Figure 4). Combining the genetic and GxE effects substantially strengthens the association signal compared to the GxE effect alone.

The *GE* test offers lower power than the *gge* test in the presence of pleiotropy except when the genetic effect is minor (*β*_g_ = 0.05) and the GxE effect is prominent (Figures 1, 2, 3). When there is no pleiotropy, the *GE* test offers lower power than the *gge* test for the first phenotype (Figure 4). The *GE* test offers lower power than the *G* test for multivariate genetic effect except when the genetic effect is small (*β*_g_ = 0.05) and the GxE effect is more potent (Figures 1, 2, 3, 4).

We now discuss the *GGE* and *G* tests’ comparative performance. First, consider the bivariate continuous phenotype. When there is no GxE effect, *β*_ge_ = 0, the *G* test produces higher power than the *GGE* test, which is expected because the GxE factor increases the degrees of freedom of the *GGE* test but does not contribute to the overall association (Figure 1). The relative power of the two tests in other scenarios depends on the magnitude of the genetic effect. When the genetic effect is small, *β*_g_ = 0.05, the *GGE* test produces higher power as the GxE effect *β*_ge_ increases (Figure 1). For *β*_g_ = 0.05 and *β*_ge_ = 0.1, the power is comparable between the two tests. For *β*_g_ = 0.05 and *β*_ge_ = 0.2, 0.3, 0.4, 0.5, the power increase of *GGE* test varies in 9 − 40% (Figure 1). As the genetic effect *β*_g_ increases, the *GGE* test requires a comparatively more substantial GxE effect to produce higher power. When *β*_g_ = 0.1, the *GGE* test results in a higher power of 6 − 11% when *β*_ge_ = 0.3, 0.4, 0.5. When *β*_g_ = 0.15, the *GGE* test leads to a very marginally higher power of 1% when *β*_ge_ = 0.3, 0.4, 0.5. We note that for *β*_g_ = 0.05, 0.1, 0.15, the *G* test offered a higher power of 1 − 7% for smaller GxE effects, *β*_ge_ = 0.1, 0.2. Because the GxE effects did not contribute adequately in the association signal to compensate for the extra two degrees of freedom in the *GGE* test. When *β*_g_ = 0.2, both approaches produce comparable power for all choices of *β*_ge_. Thus, the *GGE* test is competitive; it produces higher power than the *G* test when the genetic effect is weak, but the GxE effect is more potent. When the *GGE* test produces lower power, the power loss is limited (1 − 7%). Since both the tests suffer without pleiotropy, their relative performances remain similar to that in the presence of pleiotropy (Figure 4).

The above patterns of relative powers obtained by the *GGE* and *G* tests remain similar for a bivariate binary phenotype. The magnitudes of power differences vary marginally (Figure 2). On the other hand, the relative performance of the *GGE* test compared to the *G* test improves for a bivariate mixed phenotype. For example, when *β*_g_ = 0.15, the *GGE* test produces a higher power with an increase of 3 − 7% for *β*_ge_ = 0.3, 0.4, 0.5. However, these two tests resulted in comparable powers for the continuous phenotypes in these simulation scenarios. The power decreases of the *GGE* test than the *G* test also reduces for mixed phenotypes compared to continuous phenotypes.

Generally, the powers obtained for the continuous phenotypes were higher than those for binary or mixed phenotypes. The binary phenotype was obtained by thresholding the continuous phenotype, leading to a loss of information. The *GGE* test is advantageous when the GxE effects are more prominent than the genetic effects. The main tradeoff is whether the GxE effects across phenotypes compensate enough for the additional degrees of freedom used in the *GGE* test. When the *GGE* test loses power, the power decrease is limited. Thus, the multivariate test for combined genetic and GxE effects is a competitive approach concerning power and offers comprehensive modeling of the gene-environment interplay underlying complex phenotypes.

## 4 Real data application

We apply the methods to UK Biobank [24] data to conduct a genome-wide multivariate analysis for three lipid phenotypes: LDL cholesterol, HDL cholesterol, and triglycerides. We consider the sleep duration of the study participants as the environmental factor. Adequate sleep is a crucial lifestyle factor for well-being and mental health. Several studies communicated suggestive evidence of sleep duration being associated with lipid levels and the risk of cardiovascular diseases [34, 35, 36].

For simplicity in the analysis, we regard the individuals of White-British descent in the UK Biobank. We exclude all individuals with missing values in the phenotypes or relevant covariates, e.g., age, sex, and principal components (PCs) of genetic ancestry. After various screening, we finally considered a cohort of 353,346 individuals, comprising 163,991 males and 189,355 females. The ages of UKB participants vary in the range of 40-69 years, with a median age of 58. The lipid phenotypes showed small to moderate pair-wise correlations. HDL and LDL have a weak positive correlation of 0.1; LDL and triglycerides have a moderate positive correlation of 0.22; HDL and triglycerides have a substantial negative correlation of -0.44. We first apply log transformation to the lipid phenotype values. Then, we adjust each log-transformed phenotype for age, sex, and 20 principal components (PCs) of genetic ancestry using linear regression. We finally apply the rank-normal transformation to the adjusted residuals obtained from the phenotype-specific regressions since the methods assume normality of the phenotype distributions. The sleep duration measurements in the UK Biobank represent the average number of hours the participants sleep within 24 hours. To ensure data quality, we removed individuals with extreme sleep duration measurements, those sleeping less than two or more than 17 hours.

We implement rigorous sample quality control (QC) measures for autosomal genotype data, adhering to criteria that exclude SNPs with a minor allele frequency (MAF) below 0.01 and those significantly deviating from Hardy-Weinberg equilibrium (HWE) with a p-value less than 5 × 10^−8^. Additionally, we filter out SNPs with 10% or more missing genotypes. Ultimately, we retain 568,105 SNPs, setting the stage for comprehensive analyses. To resemble our simulation study, we focus on analyzing the GxE effect (*GE* test), genetic effect (*G* test), and combined genetic and GxE effects (*GGE* test) for a multivariate phenotype comprising the three lipids, and the Kraft’s univariate joint test for genetic and GxE effects (*gge* test) for individual lipids. We use the commonly used p-value threshold 5 × 10^−8^ adjusted for multiple testing to recognize the genome-wide significant signals.

The multivariate *GGE* and *G* tests identified 7724 and 8614 significantly associated SNPs, respectively. The univariate *gge* tests for HDL, LDL, and triglycerides found 3149, 1794, and 3036 associated SNPs, respectively. The *G* and *GGE* tests detected a common set of 7654 SNPs. Thus, the *GGE* test performed well but detected a smaller number of SNPs than the *G* test. To demonstrate the merits of the *GGE* test, we focus on finding the independent associated genetic loci that were identified by the *GGE* test but not by others, including the *G* test. The *GGE* test discovered 30 SNPs on chromosomes 2, 6, 8, 13, 14, and 19 with multivariate genetic or GxE effects, or both (Table S2), which the other competing approaches overlooked. Interestingly, these SNPs marginally missed the genome-wide p-value threshold for the multivariate genetic effect (Table S2). However, the SNPs showed significant signals of combined multivariate genetic and GxE effects (Table S2). Thus, the GxE effect complements the genetic effect for these SNPs. We note that the multivariate *GE* test also did not produce strong GxE signals for these SNPs (Table S2).

The subset of these 30 SNPs on each chromosome (Table S2) are in LD. We implement the following strategy to disentangle the independent loci identified by the *GGE* test but not by others. Using the Plink software, we apply LD pruning based on the *r*^2^ threshold of 0.2 to isolate the lead SNPs with the most vital signals. We exclude an SNP in LD with SNPs identified by the other competing approaches. Employing this strategy, we found six independent loci discovered by the *GGE* test (Table 2), which were not detected by other tests.

**Table 2:** Independent SNPs with the most vital signal of associations with lipid phenotypes discovered by the *GGE* test but missed by other competing approaches. P-values obtained by the various tests are provided. The *G* test evaluates the multivariate genetic effect. The *GE* test evaluates the multivariate GxE effect. The *GGE* test assesses the combined multivariate genetic and GxE effects. The *gge* test gauges the joint univariate genetic and GxE effects.

| SNP | CHR | BP | *G* test | *GE* test | *GGE test* | *gge* test |  |  |
| --- | --- | --- | --- | --- | --- | --- | --- | --- |
|  |  |  |  |  |  | HDL | LDL | Triglycerides |
| rs10206370 | 2 | 227368381 | 2.2E-07 | 0.001 | 5.7E-09 | 0.0002 | 0.0003 | 1.5E-06 |
| rs4947296 | 6 | 31058178 | 8.3E-07 | 0.003 | 4.7E-08 | 2.2E-05 | 0.0003 | 0.01 |
| rs3129299 | 6 | 32900787 | 5.4E-08 | 0.02 | 2.3E-08 | 9.7E-06 | 0.02 | 0.68 |
| rs2928619 | 8 | 6569927 | 3.3E-07 | 0.0004 | 3.0E-09 | 3.7E-05 | 1.4E-06 | 0.04 |
| rs111234210 | 13 | 51040660 | 8.8E-08 | 0.007 | 1.3E-08 | 1.5E-06 | 0.007 | 0.002 |
| rs887506 | 14 | 74219367 | 6.6E-08 | 0.02 | 3.1E-08 | 0.0003 | 4.0E-05 | 0.50 |

On the other hand, the *G* test discovered 47 independent loci associated with the lipids that were missed by the other approaches (Table S4). The *GGE* test overlooked these loci with marginally larger p-values compared to the genome-wide threshold (Table S4). The test used three additional degrees of freedom to evaluate the GxE effects, which did not contribute adequately to these loci.

The univariate *gge* test detected two SNPs for HDL, rs10278, and rs2289865, on chromosome 17 and 19, respectively; one SNP for triglycerides, rs888083, on chromosome 2, which were not detected by any other tests (Table S3). The two SNPs identified for HDL do not provide any distinct, independent locus, as both are in LD with other significant SNPs discovered by the *G* or the *GGE* test. However, rs888083 on chromosome 2 identified by the *gge* test for triglycerides provides an independent locus not recognized by other approaches. Thus, the univariate joint tests managed to identify only a single independent locus overlooked by others. The *GE* test for the multivariate GxE effect found no significant SNP. Thus, the multivariate *GGE* test outperformed Kraft’s univariate *gge* and multivariate *GE* tests.

Many of the identified SNPs have been reported by previous studies to be associated with the lipid phenotypes in the NHGRI-EBI GWAS catalog. For example, rs116276872 on chromosome 1 (Table S4) identified by the multivariate *G* test was communicated to be associated with LDL cholesterol levels.

However, most of the loci identified only by the multivariate *GGE* test (Table 2) are novel. rs4947296 on chromosome 6 (Table 2) detected by the *GGE* test is mapped to RNU6-1133P and C6orf15 genes. Even though the SNP itself is not associated with the lipid phenotypes, the mapped genes are associated with LDL, triglyceride, and total cholesterol levels. The other five loci (Table 2) were not reported in the NHGRI-EBI GWAS catalog. A possible reason behind the novelty of the *GGE* loci is that these do not have an individually significant genetic effect or G×E effects but a combined effect.

## 5 Discussion

We have developed an efficient testing procedure to evaluate an SNP’s joint genetic and GxE effects on a multivariate phenotype. The method extends Kraft’s joint test of the genetic and GxE effects for a univariate phenotype [1]. The approach offers a comprehensive analysis of the genetic and GxE effects for a multivariate phenotype, combining the main motives of two domains: pleiotropy and joint testing of the genetic and GxE effects. We implement the general linear hypothesis testing for a multivariate multiple linear regression of continuous phenotypes. Since a multivariate phenotype can comprise non-continuous phenotypes, we use the generalized estimating equations (GEE) technique under the seemingly unrelated regressions (SUR) framework to test for a general linear hypothesis for binary and mixed phenotypes.

Using extensive simulations, we show that our approach, the *GGE* test, outperforms the univariate joint test of the genetic and GxE effects, the *gge* test, concerning power in the presence of pleiotropy. Without pleiotropy, the *GGE* test results in a lower power than the *gge* test; however, the power loss is limited. The proposed method produces substantially higher power than testing for the multivariate GxE effects alone. Better performance of the *GGE* test than the test for multivariate genetic effect, *G* test, depends on the underlying structure of the genetic and GxE effects. The *GGE* test produces a higher power for minor genetic and significant GxE effects. The *G* test dominates when the genetic impact is more extensive. When both effects are large, the approaches lead to comparable power. Thus, the *GGE* test offers an advantage when a prominent GxE effect complements a weak genetic effect. The magnitude of power loss is limited when the *GGE* test offers lower power. We also observe a similar pattern for binary and mixed phenotypes. In the latter scenario, the relative performance of the *GGE* test improves compared to continuous phenotypes. Thus, the joint test for multivariate genetic and GxE effects is a competitive and robustly powerful approach. It comprehensively models the gene-environment interplay underlying the complex phenotypes.

We applied the methods to lipid phenotypes in the UK Biobank data, considering sleep duration as the environmental factor. The results resemble our findings in the simulation study. While the *GE* test did not discover any signal, the *gge* test identified only one independent locus for triglycerides, which other tests overlooked. The *GGE* test detected six independent loci associated with lipids, which were missed by other approaches, including the *G* test. These loci did not show significant genetic effect or GxE effect individually. Instead, they have a combined multivariate effect due to the main genetic and GxE interaction factors. The *G* test discovered a maximum number of 47 independent loci missed by other tests, including the *GGE* test. This is because the GxE effects did not adequately compensate for the additional degrees of freedom in the joint test for these loci. The *G* test performed better than the *GGE* test in the real data application to identify novel loci associated with the lipids. However, the *GGE* test offers exciting insights into GxE interaction. The GxE effects are too weak to be individually detected. The loci identified by the *GGE* test but overlooked by the *G* test are promising to investigate further because they were discovered due to some contribution from the GxE interaction factors complemented by the genetic effects. These signals are crucial from the GxE perspective because the *GGE* test performed substantially better than the other GxE approaches. Hence, we recommend the *GGE* test over the other GxE approaches considered in this paper. We put forward the test as a complementary approach to discovering genetic loci but not a substitute for the *G* test.

We point out a few limitations of our work and future directions. First, information regarding participants*ψ* medication to manage or reduce lipid levels was unavailable in the UK biobank data. Second, sleep duration was self-reported, which may be subject to measurement errors leading to power loss. Third, we did not conduct any replication study. The novel genetic loci identified by the methods are based on the discovery data alone. In future work, we aim to develop methods to adjust for the measurement errors in the environmental exposure variables while analyzing GxE interactions. Another important direction is to develop approaches to analyzing GxE interactions for a gene instead of a single SNP, which can improve the power. The transcriptome-wide association study (TWAS) framework [37] is promising in this context. In conclusion, Testing for a combined main genetic and GxE interaction effects on a multivariate phenotype is a robustly powerful approach. The developed testing procedure is technically sound and computationally efficient. We can apply the method to analyze the gene-environment interplay underlying a multivariate phenotype and understand its genetic architecture better.

## Acknowledgement

This research has been conducted using the UK Biobank Resource under Application Number 77327. We sincerely acknowledge Dr. Tanushree Haldar for helping with a primary exploration of the cloud computing platform in the UK Biobank. We thank Dr. William Gauderman for helpful discussions related to this work.

