## Supplementary Material for "A multivariate approach to joint testing of main genetic and gene-environment interaction effects"

### The estimation algorithm for GEE

Liang and Zeger [1] proposed a Fisher scoring algorithm to find the solution for the equation 8 (provided in the main text), as there is no closed-form solution. The algorithm iteratively estimates the parameters until convergence. The algorithm starts with initial estimates of the parameters obtained by ignoring the correlation structure among the phenotypes and applying generalized linear model (GLM) procedures for each equation.

After estimating the parameter $\beta$, the standardized residual or the Pearson residuals are calculated as:

$$\widehat{r}_{ij} = \frac{y_{ij} - \widehat{\mu}_{im}}{\sqrt{V(\widehat{\mu}_{ij})}}.$$

Based on these standardized residuals, the nuisance parameters $\xi$, and $\varphi$ are calculated using:

$$\widehat{\varphi}^2 = \frac{1}{n-p}\sum_{i=1}^{n}\sum_{j=1}^{m}\widehat{r}_{ij}^2$$
$$\widehat{\xi}_{jk} = \frac{1}{\widehat{\varphi}^2(n-p)}\sum_{i=1}^{n}\widehat{r}_{ij}\widehat{r}_{ik}$$

The following are the main steps:

**Initialization:**

1. Initialize the regression parameter estimates. The initial estimate of the parameter vector $\boldsymbol{\beta}$, denoted as $\boldsymbol{\beta}^{(0)}$, is obtained from the GLM estimates of the individual equations.
2. Generate the initial estimates of the parameters $\boldsymbol{\alpha} = (\xi, \varphi)'$, denoted as $\boldsymbol{\alpha}^{(0)}$, using the standardized residuals.

**Iteration Process:**

Once we have initial values, we follow the double-iteration process to estimate the parameters:

1. Given $\boldsymbol{\beta}^{(l)}$, $\boldsymbol{\alpha}^{(l)}$ is updated to $\boldsymbol{\alpha}^{(l+1)}$ using the standardized residuals based on $\boldsymbol{\beta}^{(l)}$.

2. Given $\boldsymbol{\beta}^{(l)}$ and $\boldsymbol{\alpha}^{(l+1)}$, $\hat{\boldsymbol{\beta}}$ is updated to $\boldsymbol{\beta}^{(l+1)}$ using the equation 10 (provided in the main text).

3. The above iteration is repeated until convergence.

### Wilk's lambda test statistics

Consider the following MMLR model in matrix notation for $n$ individuals' data:

$$\underset{(n\times m)}{\boldsymbol{y}} = \underset{(n\times p)}{\mathbf{X}}\underset{(p\times m)}{\mathbf{B}} + \underset{(n\times m)}{\boldsymbol{\Xi}}.$$

Here, $\underset{(n\times m)}{\boldsymbol{y}} = [\mathbf{y_1}, \cdots, \mathbf{y_m}]$ denotes the matrix of $n$ observations for $m$ phenotypes such that $\mathbf{y_j} = (y_{1j}, \cdots, y_{nj})'$ is the vector of $j^{th}$ phenotype values, $j = 1, \ldots, m$. $\underset{(n\times p)}{\mathbf{X}}$ is the design matrix with columns for $p$ regressors, including the first column of 1s corresponding to the intercept coefficient. $\underset{(p\times m)}{\mathbf{B}}$ is the matrix of regression coefficients, with one column for each phenotype. $\underset{(n\times m)}{\boldsymbol{\Xi}}$ is the matrix of error terms with columns corresponding to the phenotypes.

Let $\boldsymbol{\varepsilon_i'}$ represent the $i^{th}$ row of $\underset{(n\times m)}{\boldsymbol{\Xi}}$, then $\boldsymbol{\varepsilon_i'} \sim \mathbf{N_m}(\mathbf{0}, \boldsymbol{\Sigma})$, where $\boldsymbol{\Sigma}$ is non-singular error covariance matrix, constant across the observations. In other words, the m observations on the $j^{th}$ sample unit have covariance matrix $\Sigma$ ), but the errors for different sample units are not correlated; $\varepsilon_i'$ and $\varepsilon_j'$ are independent for $i \neq j$. $X$ is fixed or independent of $\underset{(n\times m)}{\boldsymbol{\Xi}}$. Further, we can write $vec(\boldsymbol{\Xi}) \sim \mathbf{N_{nm}}(\mathbf{0}, \mathbf{I_n} \otimes \boldsymbol{\Sigma})$, where $vec(\boldsymbol{\Xi})$ is the error matrix row-wise into a vector, $\mathbf{I_n}$ is the identity matrix of order n, and $\otimes$ is the Kronecker-Product operator.

The maximum likelihood estimator of $\mathbf{B}$ in the multivariate linear model is

$$\hat{\mathbf{B}} = (\mathbf{X}'\mathbf{X})^{-\mathbf{1}}\mathbf{X}'\mathbf{Y}.$$

The decomposition into sums of squares and cross products:

$$\underbrace{\mathbf{Y}'\mathbf{Y}}_{\text{TotSSCP}} = \underbrace{\widehat{\mathbf{Y}}'\widehat{\mathbf{Y}}}_{\text{RegSSCP}} + \underbrace{\widehat{\boldsymbol{\Xi}}'\widehat{\boldsymbol{\Xi}}}_{\text{ResSSCP}}.$$

Where TotSSCP stands for the total sums of squares and cross-products matrix, RegSSCP stands for the regression sums of squares and cross-products matrix, and ResSSCP stands for the residual sums of squares and cross-products matrix.

Suppose that we want to test the linear hypothesis

$$H_0 : \underset{(q\times p)}{\mathbf{L}}\underset{(p\times m)}{\mathbf{B}} = \underset{(q\times m)}{\mathbf{C}}, q \leq p.$$

where $\mathbf{L}$ is a hypothesis matrix of full row-rank $q \leq p$, and the right-hand-side matrix $\mathbf{C}$ consists of constants, usually 0s. Then the SSCP matrix for the hypothesis is defined as [38]:

$$\mathbf{SSCP}_H = \left(\widehat{\mathbf{B}}'\mathbf{L}' - \mathbf{C}'\right)\left[\mathbf{L}\left(\mathbf{X}'\mathbf{X}\right)^{-1}\mathbf{L}'\right]^{-1}(\mathbf{L}\widehat{\mathbf{B}} - \mathbf{C}).$$

Where $\mathbf{SSCP}_H$ represent the incremental SSCP matrix for a hypothesis, that is, the difference between $\mathbf{SSP_{reg}}$ for the model unrestricted by the hypothesis and $\mathbf{SSCP_{reg}}$ for model on which the hypothesis is imposed. Multivariate tests for the hypothesis are based on the $m$ eigenvalues $\lambda_j$ of $\mathbf{SSP}_H\mathbf{SSCP}_R^{-1}$ (the hypothesis SSCP matrix "divided by" the residual SSCP matrix), that is, the values of $\lambda$ for which

$$\det\left(\mathbf{SSCP}_H\mathbf{SSCP}_R^{-1} - \lambda\mathbf{I}_m\right) = 0.$$

Based on these eigenvalues, the Wilk's lambda test statistic is defined as:

$$\text{Wilks's Lambda, } \Lambda = \prod_{j=1}^{m} \frac{1}{1+\lambda_j}.$$

The tests generally apply to all linear hypotheses.For different hypothesis matrices $\mathbf{L}$, different linear hypotheses can be tested.

Table S1: Estimated type I error rates for univariate joint tests for various choices of the null hypotheses. Subscripts 1 and 2 represent the first and second phenotypes, respectively. In the mixed phenotype case, $gge$c represents the continuous phenotype, and $gge$b represents the binary phenotype.

| | | **Tests for univariate continuous phenotypes** | | **Tests for univariate mixed phenotypes** | | **Tests for univariate binary phenotypes** | |
|---|---|---|---|---|---|---|---|
| MAF | P(E=1) | $gge_1$ | $gge_2$ | $gge_c$ | $gge_b$ | $gge_1$ | $gge_2$ |
| 0.1 | 0.1 | 0.06 | 0.05 | 0.04 | 0.05 | 0.06 | 0.05 |
| 0.1 | 0.25 | 0.04 | 0.05 | 0.05 | 0.05 | 0.05 | 0.05 |
| 0.25 | 0.1 | 0.05 | 0.05 | 0.05 | 0.05 | 0.05 | 0.06 |
| 0.25 | 0.25 | 0.05 | 0.04 | 0.05 | 0.04 | 0.06 | 0.04 |

Table S2: SNPs discovered by multivariate GGE test only. P-values obtained by the various tests are provided. The *G* test evaluates the multivariate genetic effect. The *GE* test evaluates the multivariate GxE effect. The *GGE* test assesses the combined multivariate genetic and GxE effects. The *gge* test gauges the joint univariate genetic and GxE effects.

| | | | | | | *gge* test | | |
|---|---|---|---|---|---|---|---|---|
| CHR | SNP | BP | *G* test | *GE* test | *GGE test* | HDL | LDL | Triglycerides |
| 2 | rs10206370 | 227368381 | 2.2e-07 | 0.001 | 5.7e-09 | 0.0002 | 0.0003 | 1.5e-06 |
| 6 | rs29273 | 29600000 | 6.1e-08 | 0.014 | 1.8e-08 | 1.0e-06 | 0.1 | 0.023 |
| 6 | rs29231 | 29600000 | 1.4e-07 | 0.012 | 3.5e-08 | 1.1e-05 | 0.067 | 0.015 |
| 6 | rs3129090 | 29700000 | 9.7e-08 | 0.011 | 2.2e-08 | 1.8e-06 | 0.12 | 0.032 |
| 6 | rs1736946 | 29800000 | 5.5e-08 | 0.028 | 3.4e-08 | 4.6e-07 | 0.19 | 0.69 |
| 6 | rs2844725 | 30500000 | 1.3e-07 | 0.018 | 4.7e-08 | 0.97 | 3.2e-06 | 0.22 |
| 6 | rs2844724 | 30500000 | 5.3e-08 | 0.013 | 1.5e-08 | 0.97 | 1.3e-06 | 0.24 |
| 6 | rs2844723 | 30500000 | 1.8e-07 | 0.013 | 4.53e-08 | 0.94 | 5.0e-06 | 0.19 |
| 6 | rs2516662 | 30500000 | 5.8e-08 | 0.022 | 2.7e-08 | 0.99 | 1.6e-06 | 0.24 |
| 6 | rs2844720 | 30500000 | 8.7e-08 | 0.016 | 3.0e-08 | 0.99 | 2.3e-06 | 0.21 |
| 6 | rs2023609 | 30500000 | 8.1e-08 | 0.014 | 2.4e-08 | 0.99 | 2.3e-06 | 0.19 |
| 6 | rs996589 | 30500000 | 6.5e-08 | 0.022 | 3.1e-08 | 0.97 | 1.98e-06 | 0.19 |
| 6 | rs2844718 | 30500000 | 9.0e-08 | 0.01 | 1.9e-08 | 0.92 | 1.8e-06 | 0.26 |
| 6 | rs975195 | 30500000 | 1.2e-07 | 0.01 | 2.9e-08 | 0.99 | 2.7e-06 | 0.19 |
| 6 | rs9366764 | 30979793 | 2.8e-07 | 0.002 | 1.1e-08 | 5.7e-05 | 3.4e-05 | 0.52 |
| 6 | rs9380205 | 30981361 | 3.5e-07 | 0.002 | 1.4e-08 | 9.1e-05 | 2.8e-05 | 0.49 |
| 6 | rs9391701 | 30983263 | 5.2e-07 | 0.002 | 1.9e-08 | 9.5e-05 | 3.2e-05 | 0.53 |
| 6 | rs3871466 | 30983683 | 1.9e-07 | 0.002 | 7.9e-09 | 0.0001 | 1.9e-05 | 0.4 |
| 6 | rs3869096 | 30984404 | 2.8e-07 | 0.002 | 1.2e-08 | 7.6e-05 | 3.2e-05 | 0.49 |
| 6 | rs9295942 | 30987010 | 2.2e-07 | 0.002 | 8.2e-09 | 7.0e-05 | 2.7e-05 | 0.47 |
| 6 | rs9380215 | 31049655 | 7.1e-07 | 0.002 | 4.9e-08 | 3.5e-05 | 0.0003 | 0.009 |

| | | | | | | | | |
|---|---|---|---|---|---|---|---|---|
| 6 | rs4947296 | 31100000 | 8.3e-07 | 0.003 | 4.7e-08 | 2.2e-05 | 0.0003 | 0.01 |
| 6 | rs114963023 | 31200000 | 2.7e-07 | 0.006 | 3.2e-08 | 0.0001 | 0.0002 | 0.004 |
| 6 | rs7775228 | 32700000 | 9.4e-08 | 0.006 | 1.2e-08 | 0.004 | 0.003 | 0.007 |
| 6 | rs3129299 | 32900000 | 5.4e-08 | 0.02 | 2.3e-08 | 9.6e-06 | 0.02 | 0.68 |
| 8 | rs2928579 | 6610000 | 4.5e-06 | 0.0005 | 4.3e-08 | 8.3e-06 | 8.5e-05 | 0.01 |
| 8 | rs11136344 | 145059425 | 6.9e-07 | 0.003 | 4.5e-08 | 0.4 | 1.3e-07 | 0.18 |
| 13 | rs111234210 | 51040660 | 8.8e-08 | 0.007 | 1.3e-08 | 1.5e-06 | 0.007 | 0.002 |
| 14 | rs887506 | 74200000 | 6.6e-08 | 0.02 | 3.1e-08 | 0.0003 | 4.0e-05 | 0.5 |
| 19 | rs73057960 | 50100000 | 1.9e-07 | 0.004 | 1.4e-08 | 0.05 | 0.005 | 0.0007 |

Table S3: Independent SNPs discovered in univariate gge tests. P-values obtained by the various tests are provided. The *G* test evaluates the multivariate genetic effect. The *GE* test evaluates the multivariate GxE effect. The *GGE* test assesses the combined multivariate genetic and GxE effects. The *gge* test gauges the joint univariate genetic and GxE effects.

| | | | | | | *gge* test | | |
|---|---|---|---|---|---|---|---|---|
| SNP | CHR | BP | *G* test | *GE* test | *GGE test* | HDL | LDL | Triglycerides |
| rs10278 | 17 | 46939658 | 5.2E-07 | 0.11 | 1.1E-06 | 4.2E-08 | 0.7 | 0.0003 |
| rs2289865 | 19 | 4013322 | 7.7E-07 | 0.04 | 5.1E-07 | 2.5E-08 | 0.7 | 0.06 |
| rs888083 | 2 | 37117073 | 1.2E-06 | 0.11 | 2.5E-06 | 0.006 | 0.2 | 4.3E-08 |

Table S4: SNPs discovered by multivariate G test only. P-values obtained by the various tests are provided. The *G* test evaluates the multivariate genetic effect. The *GE* test evaluates the multivariate GxE effect. The *GGE* test assesses the combined multivariate genetic and GxE effects. The *gge* test gauges the joint univariate genetic and GxE effects.

| | | | | | | *gge* test | | |
|---|---|---|---|---|---|---|---|---|
| CHR | SNP | BP | *G* test | *GE* test | *GGE test* | HDL | LDL | Triglycerides |
| 1 | rs116276872 | 110068291 | 2.03e-08 | 0.297 | 1.55e-07 | 0.0214 | 0.000983 | 0.137 |
| 1 | rs11102516 | 113161350 | 4.85e-08 | 0.631 | 8.5e-07 | 0.366 | 0.00012 | 0.000103 |
| 2 | rs77998815 | 119767026 | 2.12e-08 | 0.825 | 5.74e-07 | 0.612 | 0.921 | 7.64e-07 |
| 2 | rs72911133 | 174772660 | 1.47e-09 | 0.86 | 5.11e-08 | 0.0254 | 0.28 | 0.0139 |
| 3 | rs62246446 | 47185650 | 1.3e-08 | 0.424 | 1.54e-07 | 5.36e-07 | 0.214 | 0.553 |
| 4 | rs13116684 | 48037926 | 9.2e-09 | 0.367 | 9.43e-08 | 0.0522 | 0.0355 | 0.0492 |
| 4 | rs1714013 | 57942167 | 9.81e-09 | 0.654 | 2.02e-07 | 0.0115 | 0.000231 | 0.609 |
| 5 | rs2112161 | 72919887 | 8.07e-09 | 0.642 | 1.64e-07 | 0.00013 | 0.883 | 0.123 |
| 5 | rs4705873 | 132267167 | 1.95e-08 | 0.316 | 1.61e-07 | 0.427 | 0.96 | 2.78e-06 |

| | | | | | | | | |
|---|---|---|---|---|---|---|---|---|
| 5 | rs6873053 | 156376703 | 5.59e-09 | 0.982 | 2.32e-07 | 0.043 | 1.44e-06 | 0.00575 |
| 6 | rs7756992 | 20679709 | 3.71e-08 | 0.0936 | 7.93e-08 | 0.539 | 0.011 | 2.3e-05 |
| 6 | rs198820 | 26124243 | 1.53e-08 | 0.287 | 1.15e-07 | 0.2 | 0.000106 | 0.15 |
| 6 | rs9461366 | 27310533 | 3.56e-08 | 0.723 | 7.64e-07 | 0.0977 | 0.172 | 0.0161 |
| 6 | rs9400897 | 116599665 | 3.26e-08 | 0.787 | 7.95e-07 | 0.00325 | 6.68e-06 | 0.128 |
| 6 | rs12214416 | 160910517 | 4.9e-09 | 0.957 | 1.92e-07 | 0.0824 | 0.0437 | 0.0542 |
| 6 | rs41272114 | 161006077 | 6.52e-09 | 0.712 | 1.55e-07 | 3.15e-05 | 0.0103 | 0.738 |
| 7 | rs62451583 | 36169097 | 3.69e-08 | 0.121 | 1.03e-07 | 0.000485 | 0.0292 | 0.243 |
| 7 | rs6462657 | 36201713 | 6.35e-09 | 0.432 | 8.05e-08 | 2.14e-06 | 0.349 | 0.593 |
| 8 | rs330052 | 9088245 | 2.32e-08 | 0.809 | 6.03e-07 | 0.448 | 0.138 | 2.86e-05 |
| 8 | rs79680096 | 10166265 | 4.18e-08 | 0.135 | 1.3e-07 | 0.11 | 0.316 | 5.97e-06 |
| 8 | rs76245571 | 11228757 | 1.82e-08 | 0.169 | 7.58e-08 | 0.204 | 0.0646 | 0.00102 |
| 8 | rs11782559 | 11804882 | 2.08e-09 | 0.7 | 5.21e-08 | 0.0202 | 0.0473 | 0.0777 |
| 8 | rs75215518 | 11841500 | 6.98e-09 | 0.733 | 1.72e-07 | 0.109 | 0.687 | 0.000622 |
| 9 | Affx-33687749 | 33113121 | 2.73e-08 | 0.433 | 3.16e-07 | 0.00392 | 0.737 | 0.0344 |
| 11 | rs77498573 | 48350646 | 2.64e-08 | 0.521 | 3.81e-07 | 0.947 | 0.786 | 7.63e-06 |
| 11 | rs500161 | 65695438 | 2.97e-08 | 0.889 | 8.87e-07 | 0.917 | 4.06e-05 | 0.109 |
| 11 | rs75656082 | 75807530 | 2.9e-08 | 0.156 | 1.08e-07 | 3.41e-05 | 0.245 | 0.386 |
| 11 | rs8177376 | 126163612 | 4.77e-08 | 0.356 | 4.23e-07 | 0.00658 | 0.000655 | 0.954 |
| 11 | rs34271700 | 126328903 | 5.44e-09 | 0.905 | 1.89e-07 | 1.57e-06 | 0.0904 | 0.51 |
| 12 | rs35117 | 46242195 | 2.43e-08 | 0.151 | 8.82e-08 | 0.472 | 0.726 | 9.93e-06 |
| 13 | rs7318105 | 95271934 | 2.61e-09 | 0.794 | 7.69e-08 | 0.0423 | 0.208 | 0.0165 |
| 15 | rs72770737 | 101869604 | 1.07e-08 | 0.539 | 1.72e-07 | 0.368 | 0.0379 | 0.00021 |
| 15 | rs7171582 | 102051266 | 3.01e-08 | 0.192 | 1.4e-07 | 0.00837 | 0.233 | 0.0198 |
| 16 | rs7186852 | 30635659 | 4.62e-08 | 0.246 | 2.71e-07 | 0.193 | 1.54e-06 | 0.000988 |
| 16 | rs117278268 | 55981520 | 2.4e-09 | 0.708 | 6.05e-08 | 7.36e-06 | 0.362 | 0.775 |
| 16 | rs592196 | 56575062 | 3.53e-08 | 0.639 | 6.45e-07 | 1.04e-06 | 0.293 | 0.222 |

| | | | | | | | | |
|---|---|---|---|---|---|---|---|---|
| 16 | rs12933525 | 56735443 | 2.66e-08 | 0.9 | 8.18e-07 | 1.28e-06 | 0.338 | 0.726 |
| 16 | rs11075743 | 69903803 | 1.86e-08 | 0.178 | 8.17e-08 | 0.568 | 0.179 | 4.55e-05 |
| 17 | rs117288663 | 26749941 | 3.28e-08 | 0.378 | 3.2e-07 | 0.137 | 0.00378 | 0.0803 |
| 17 | rs883541 | 66449122 | 2.01e-08 | 0.828 | 5.49e-07 | 0.825 | 0.00999 | 0.00986 |
| 18 | rs4464148 | 46459032 | 8.39e-09 | 0.766 | 2.17e-07 | 0.00436 | 0.503 | 0.046 |
| 19 | rs11545166 | 10671894 | 4.71e-08 | 0.709 | 9.64e-07 | 0.877 | 5.91e-06 | 0.248 |
| 19 | rs17714646 | 45042495 | 3.64e-09 | 0.989 | 1.59e-07 | 0.0944 | 2.89e-05 | 0.912 |
| 19 | rs2357100 | 45099240 | 8.55e-09 | 0.762 | 2.19e-07 | 6.37e-05 | 0.0534 | 0.951 |
| 19 | rs57579470 | 46005824 | 1.34e-08 | 0.919 | 4.49e-07 | 0.899 | 2.48e-05 | 0.147 |
| 19 | rs73048457 | 46089055 | 4.38e-09 | 0.604 | 8.6e-08 | 0.569 | 2.29e-05 | 0.192 |
| 20 | rs117590445 | 61321110 | 6.58e-09 | 0.669 | 1.43e-07 | 0.138 | 0.00448 | 0.0925 |
